## Supplementary Material for "Spontaneous pauses in firing of external pallidum neurons are associated with exploratory behavior"

<u>**Extended Data Figures**</u>

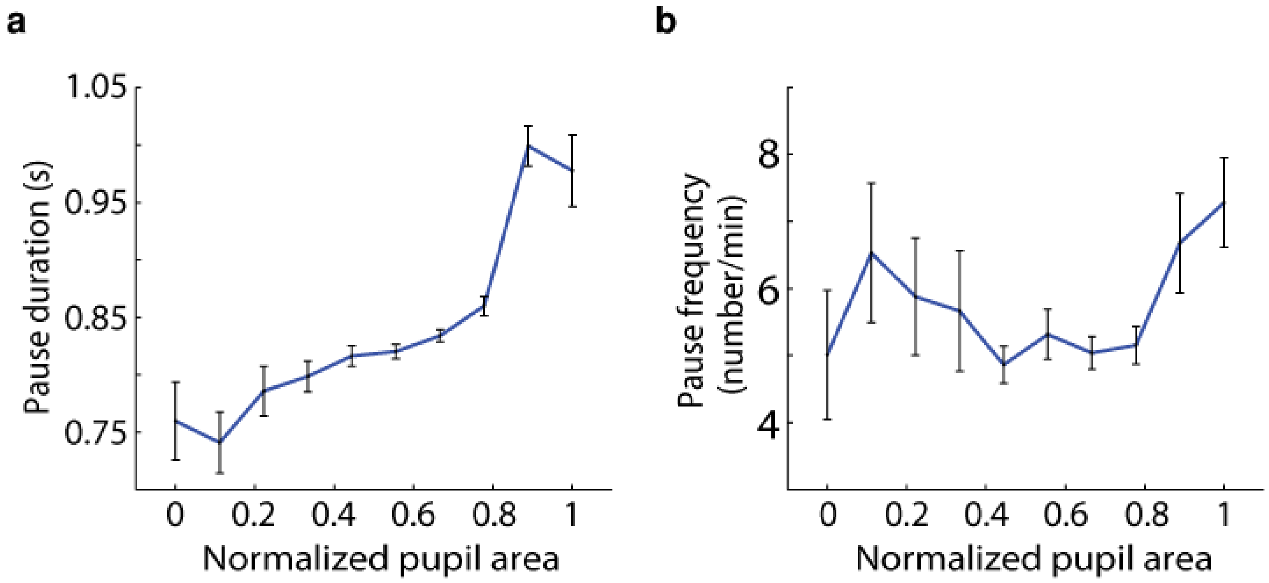


**Extended Data Figure 1. Dilated pupils are associated with both longer and more frequent pauses.**

**(a)** Average pause duration as a function of pupil area at the time of pause initiation (*n* = 24,968 pauses). Abscissa, normalized pupil area (values between 0-0.1, 0.1-0.2,..., 0.8-0.9, 0.9-1); ordinate, pause duration in seconds. Error bars represent s.e.m. Only pauses that corresponded to real-valued pupil area (not NaN) were included in the analysis. **(b)** Average pause frequency as a function of pupil area at the time of pause initiation (*n* = 579 neurons). Abscissa, normalized pupil area; ordinate, pause frequency in numbers per minute. Error bars represent s.e.m. Only pauses that corresponded to real-valued pupil area (not NaN) were included in the analysis.

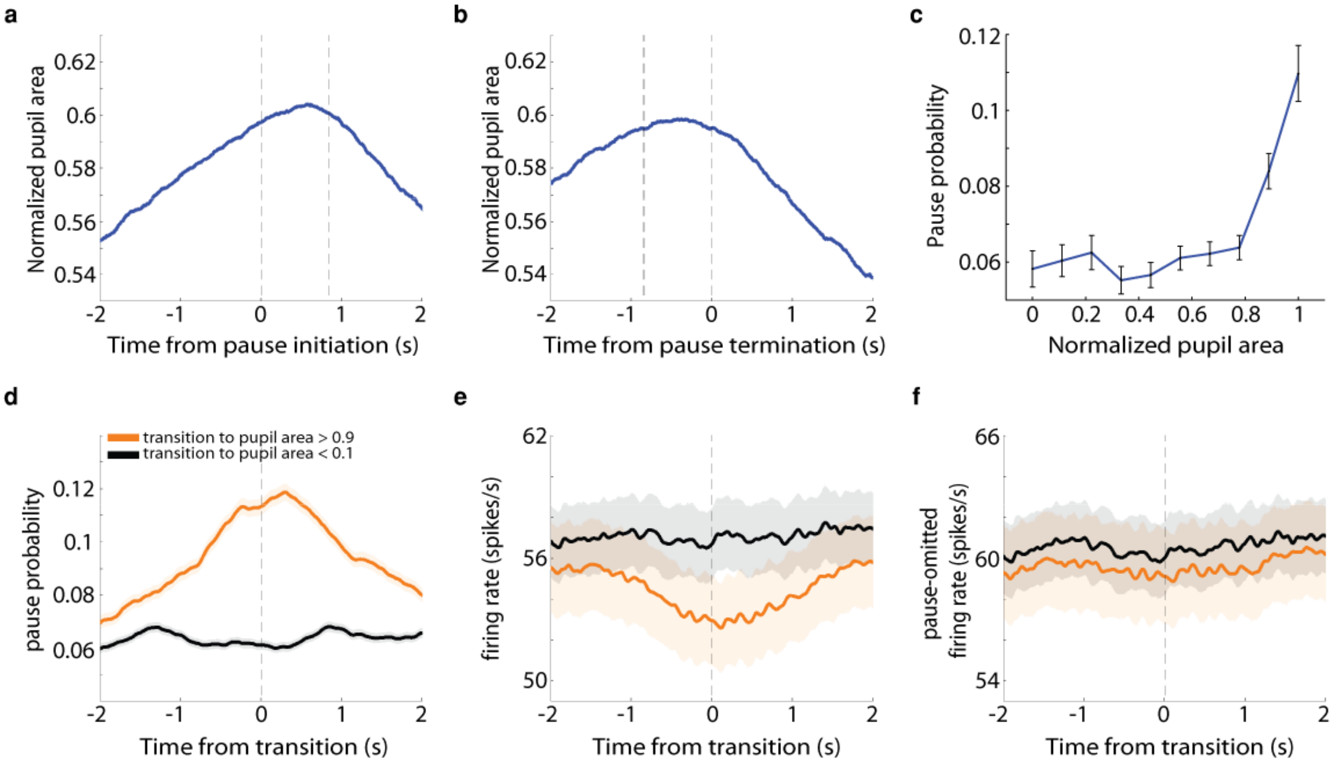


**Extended Data Figure 2. GPe pause probability is increased when the pupils are dilated, even when considering only periods of time when no saccades were made.**

**(a)** Average pupil area around pauses (*n* = 31,247 pauses). Abscissa, time (−2 to 2 s); ordinate, pupil area (scaled using min-max normalization). The vertical dashed lines at t = 0 and t = ~0.85 s indicate the time of pause initiation and mean pause termination, respectively. Shaded regions represent s.e.m. Only data from time periods when no saccades were made were considered for analysis. **(b)** Same as (a), but the vertical dashed lines at t = 0 and t = ~-0.85 s indicate the time of pause termination and mean pause initiation, respectively. **(c)** Average population pause probability as a function of normalized pupil area (*n* = 579 neurons). For each neuron and for each range of pupil area values (0-0.1, 0.1-0.2,…, 0.8-0.9, 0.9-1), the total time the neuron paused was divided by the overall time the pupil was in a particular area range. Error bars represent s.e.m. Only data from time periods when no saccades were made were considered for analysis. **(d)** Average population dynamics of pause probability for different normalized pupil areas. Abscissa, time (−2 to 2 s), zero is the time when the normalized pupil area crossed to values < 0.1 (black curve; *n* = 17,507 transitions) and to values > 0.9 (orange curve; *n* = 10,605 transitions); ordinate, pause probability. The pause probability was computed using 1-ms bins and smoothed with a Gaussian window with a s.d. of 20 ms. Shaded regions represent s.e.m. Only data from time periods when no saccades were made were considered for analysis. **(e)** Average population dynamics of discharge rate for different normalized pupil areas. Abscissa, time (−2 to 2 s), zero is the time when the normalized pupil area crossed to values < 0.1 (black curve; *n* = 17,507 transitions) and to values > 0.9 (orange curve; *n* = 10,605 transitions); ordinate, firing rate in Hz. Same conventions as in (d). **(f)** Average population dynamics of discharge rate after removal of pause-containing segments. Same conventions as in (e).

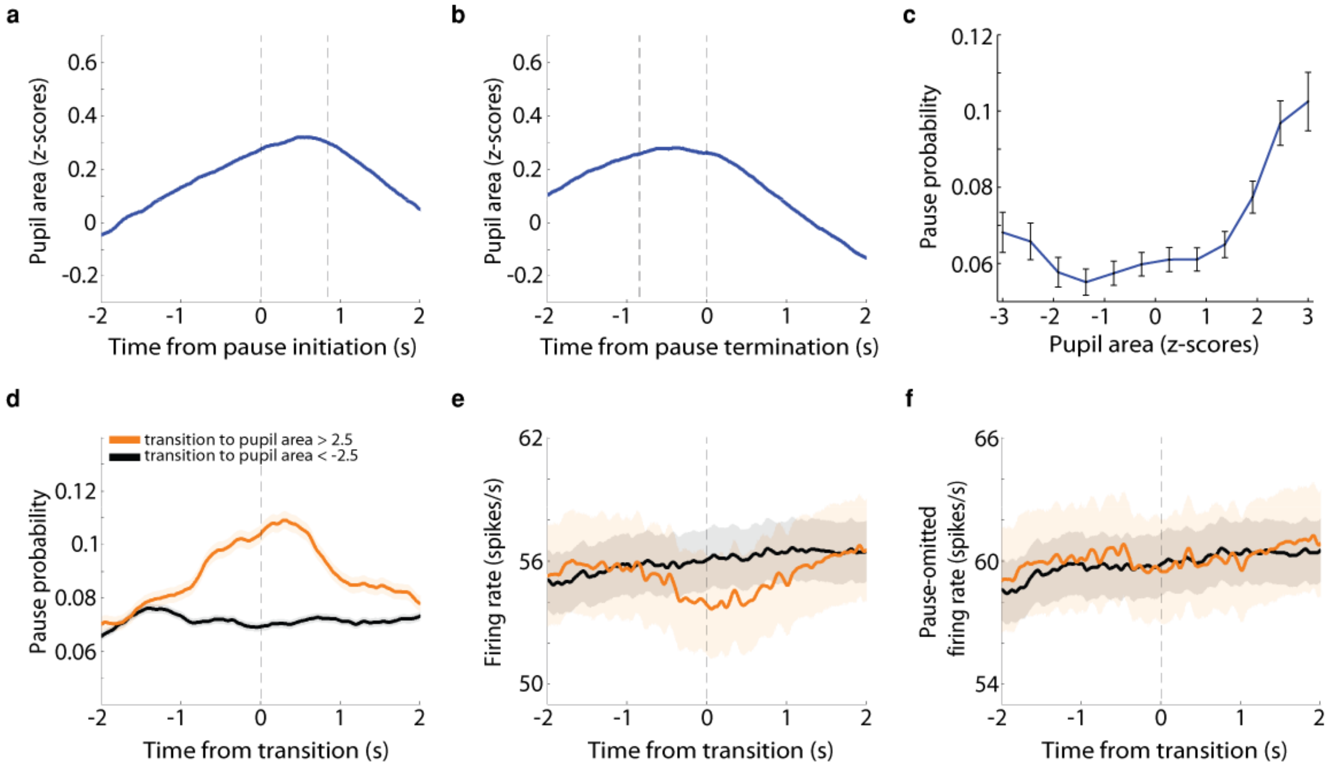


**Extended Data Figure 3. GPe pause probability is increased when the pupils are dilated, regardless of the normalization method used.**
**(a)** Average pupil area around pauses (*n* = 31,247 pauses). Abscissa, time (−2 to 2 s); ordinate, pupil area (normalized using z-scores). The vertical dashed lines at t = 0 and t = ~0.85 s indicate the time of pause initiation and mean pause termination, respectively. Shaded regions represent s.e.m. **(b)** Same as (a), but the vertical dashed lines at t = 0 and t = ~-0.85 s indicate the time of pause termination and mean pause initiation, respectively. **(c)** Average population pause probability as a function of *z*-score-normalized pupil area (*n* = 579 neurons). For each neuron and for each range of pupil area values (-3 to -2.5, -2.5 to -2, -2 to -1.5,…, 2 to 2.5, 2.5 to 3), the total time the neuron paused was divided by the overall time the pupil was in a particular area range. Error bars represent s.e.m. **(d)** Average population dynamics of pause probability for different normalized pupil areas. Abscissa, time (−2 to 2 s), zero is the time when the normalized pupil area crossed to *z*-score < -2.5 (black curve; *n* = 23,971 transitions) and to *z*-score > 2.5 (orange curve; *n* = 8,735 transitions); ordinate, pause probability. The pause probability was computed using 1-ms bins and smoothed with a Gaussian window with a s.d. of 20 ms. Shaded regions represent s.e.m. **(e)** Average population dynamics of discharge rate for different normalized pupil areas. Abscissa, time (−2 to 2 s), zero is the time when the normalized pupil area crossed to *z*-score < -2.5 (black curve; *n* = 23,971 transitions) and to *z*-score > 2.5 (orange curve; *n* = 8,735 transitions); ordinate, firing rate in Hz. Same conventions as in (d). **(f)** Average population dynamics of discharge rate after removal of pause-containing segments. Same conventions as in (e).

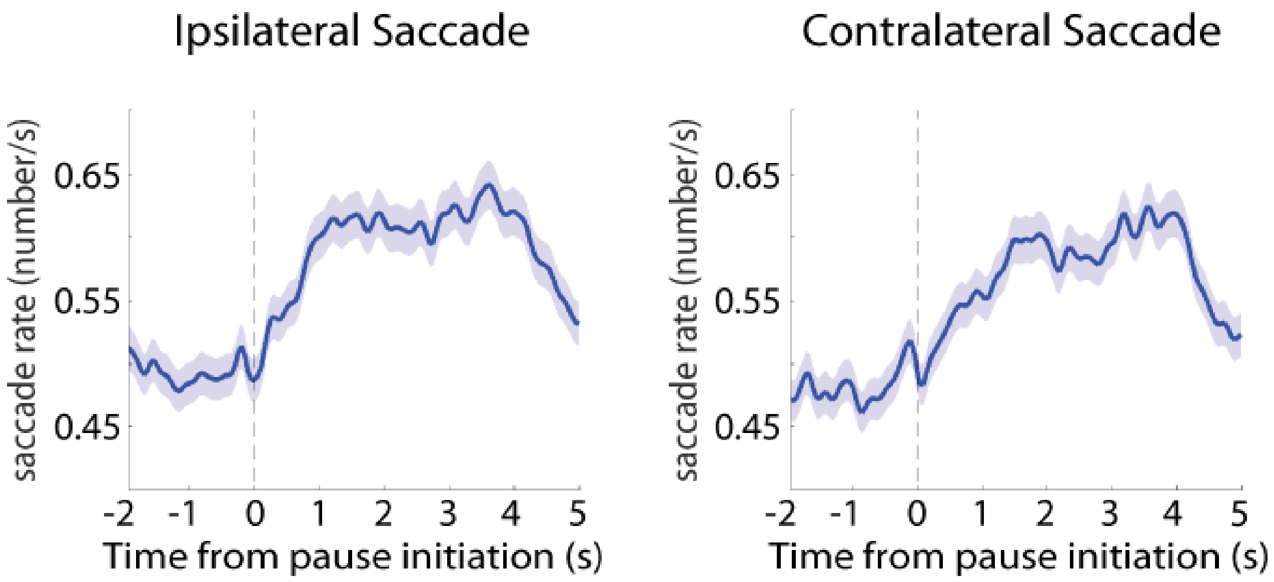


**Extended Data Figure 4. Saccades to the contralateral visual field are not more abundant around GPe pauses than saccades to the ipsilateral visual field.**

Average saccade rate to the ipsilateral and contralateral visual fields around pauses (*n* = 33,798 pauses). Abscissa, time (−2 to 5 s, time zero indicates pause initiation); ordinate, saccade rate in Hz. Saccade rate was computed using 50-ms bins and smoothed with a Gaussian window with a s.d. of 1 ms. Shaded regions represent s.e.m.
